# Drug mode of action and resource constraints modulate antimicrobial resistance evolution

**DOI:** 10.1101/2023.08.29.555413

**Authors:** Oscar Delaney, Andrew D. Letten, Jan Engelstädter

## Abstract

One of the key characteristics of an antibiotic drug is its mode of action: bacteriostatic or bactericidal. The effect of drug mode of action on the evolution of resistance has been surprisingly underinvestigated to date. We present a theoretical model comparing the efficacy of bacteriostatic and bactericidal drugs, and drugs of intermediate type, at preventing the evolutionary rescue of an initially susceptible bacterial population in a patient. Our results suggest that in resource-abundant environments bacteriostatic drugs are best, as they constrain cell divisions and thus cause fewer resistance mutations to occur. When multiple drugs are employed, using one bacteriostatic and one bactericidal drug is usually optimal, because the cell division rate cannot fall below zero, so there are diminishing returns to bacteriostatic activity from two drugs. We also provide a web-based simulation engine for other researchers to intuitively explore related dynamics without requiring programming expertise.

## 1 Introduction

Antimicrobial resistance (AMR) is a large and growing problem: in 2019 over 1 million people are estimated to have died due to AMR infections [1]. Therefore, one of the key goals of designing antimicrobial treatment regimens must be to minimize the probability that resistance develops, alongside striving to rapidly clear the patient’s infection and avoid excessive toxicity.

Antimicrobials are often classified by mode of action: bactericidal drugs are those that increase mortality, and bacteriostatic drugs inhibit growth [2, 3]. This division is not binary — drugs in reality often exhibit both bactericidal and bacteriostatic functionality, or may have different characteristics depending on the dosage and the type of bacteria [4, 5]. However, the mode of action continuum remains a useful conceptual framework. Specifically, when designing multidrug treatment regimens to minimise resistance evolution, it is valuable to consider what pairings of bacteriostatic and bactericidal antibiotics would be most appropriate.

Two prominent multidrug treatment strategies are combination therapy and cycling therapy. In combination therapy, two (or more) drugs are applied simultaneously, the rationale being that if a mutant with resistance to one of the drugs arises, it will not be able to establish itself because it would still be susceptible to the second drug [6]. Cycling therapy involves applying one drug at a time, but rotating through two (or more) drugs, such that even if resistance to the drug currently being applied develops, when the second drug is administered the resistant mutants will then be selected against [7, 8, 9]. For each of single-drug, cycling, and combination therapy, we used a mathematical model and computer simulation to investigate which modes of action are most effective in preventing resistance evolution while still eliminating the infection. We find that under resource-abundant conditions, bacteriostatic drugs are better at preventing resistance evolution, but that this effect reverses as resource limitations become more binding.

## 2 Methods

Our mathematical model is based on the one formulated by Nyhoegen and Uecker [16]. All parameters are described in Table 1, with supporting literature where appropriate. Like Nyhoegen and Uecker we included two drugs, *A* and *B*, with concentrations *C*_*A*_(*t*) and *C*_*B*_(*t*) respectively, and four bacterial genotypes: resistant to neither drug, *A* only, *B* only, or both *A* and *B*. The state vector of population sizes was defined as

**Table 1:** All simulation parameters with their default values and sources where applicable. When the parameter varies by strain, values are given in the order [*S, A, B, AB*].

| Parameter | Description | Default value | Reference |
| --- | --- | --- | --- |
| $R(0)$ | Initial resource concentration | $10^{11} \text{ mg L}^{-1}$ | - |
| $\mathbf{N}(0)$ | Initial population sizes | $[5 \cdot 10^8, 0, 0, 0]$ | - |
| $C_j(0)$ | Drug $j$ influx concentration | $3 \cdot z_{S,j}$ | - |
| $\omega$ | Time between drug doses | 10 h | - |
| $\mu_i$ | Maximum growth rate for strain $i$ | $1 \text{ h}^{-1}$ | [10, 11, 12] |
| $\delta_i$ | Intrinsic death rate | $0.4 \text{ h}^{-1}$ | - |
| $\theta_j$ | Maximum drug $j$ induced death rate | $1 \text{ h}^{-1}$ | - |
| $\phi_j$ | Maximum bacteriostatic effect of drug $j$ | 1 | - |
| $z_{i,A}$ | Drug $A$ concentration at half-max effect | $[1, 28, 1, 28] \mu\text{g L}^{-1}$ | [11, 12] |
| $z_{i,B}$ | Drug $B$ concentration at half-max effect | $[1, 1, 28, 28] \mu\text{g L}^{-1}$ | [11, 12] |
| $\beta_j$ | Hill function shape parameter for drug $j$ | 1 | [13] |
| $m_j$ | Mutation rate for $j$-resistance per replication | $10^{-9}$ | [14, 15] |
| $\gamma_j$ | Rate of drug $j$ elimination | $0.1 \text{ h}^{-1}$ | [16] |
| $K$ | $R$ at half-maximal growth rate | $10^8 \text{ mg L}^{-1}$ | - |
| $\alpha$ | Resources used per replication | $1 \text{ mg L}^{-1}$ | - |
| $R_{\text{in}}$ | Resource supply rate | $10^8 \text{ mg L}^{-1} \text{ h}^{-1}$ | - |

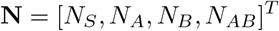

We extended Nyhoegen and Uecker’s logistic-growth model to be resource explicit, by introducing another state variable for the concentration *R* of a single growth-limiting resource, and using the Monod equation [17] to model the growth of strain *i* in the absence of drugs as:

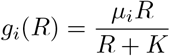

We followed Nyhoegen and Uecker’s pharmacokinetic approach of letting the drug concentrations decay exponentially over time, however we diverged on the pharmacodynamics. While their analysis was limited to bactericidal drugs only, we modelled any combination of bactericidal/static activity. Specifically, we first normalised the effective drug concentration for bacterial strain *i* and drug *j* onto the [0, 1) interval using the sigmoid E_max_ model [18]:

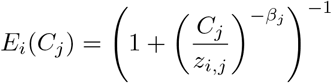

Using *θ* and *ϕ* to represent the bactericidal and bacteriostatic activity of each drug, respectively, we defined the drug- and resource-dependent growth rate and death rate row vectors as:

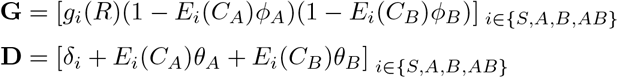

Ignoring back-mutations, the probability that a replication event in strain *i* leads to a new cell in strain *i*^′^ can be represented by:

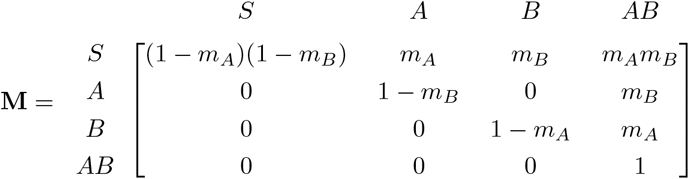

We defined our system of ODEs, using ◦ for element-wise Hadamard multiplication, as:

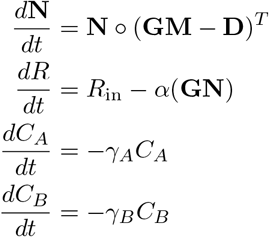

Finally, starting at *t* = 0, drugs are added to the system every *w* hours, either just *A* in monotherapy, both *A* and *B* in combination therapy, or *A* and *B* alternating in cycling therapy. Along with the initial values of all seven state variables, this fully defines the mathematical model. For the computational implementation, in order to more realistically model the uncertainty inherent in growth and mutation, we used the Stochastic Simulation Algorithm to evolve the system over time, with separate birth and death events [19]. To make the system computationally feasible, we used tau-leaping to perform many transitions in one step [20, 21]. All simulations were performed using R [22].

## 3 Results

We considered an evolutionary rescue scenario where a patient has some large susceptible bacterial population at *t* = 0 when treatment begins, and either resistance will develop and the bacterial population survives, or resistance fails to emerge and the bacterial population goes towards extinction. We first considered a simple scenario with abundant resources and using just drug *A*, which had a fixed total effect but varied as to how bactericidal or bacteriostatic it was, that is *θ*_*A*_ + *ϕ*_*A*_ = 1. Example population dynamics are shown in Figure 1A. Because the total activity of the drug was fixed, the decline in the susceptible bacterial population was equivalent across all drug *A* modes of action (not shown). The probability that the population goes extinct increases as the mode of action of the drug changes from fully bactericidal to fully bacteriostatic (Figure 1B). This is because in our model mutation is coupled to replication, and so if a bacteriostatic drug causes fewer cell divisions to occur, there will be concomitantly fewer chances for resistance to emerge.

**Figure 1:**
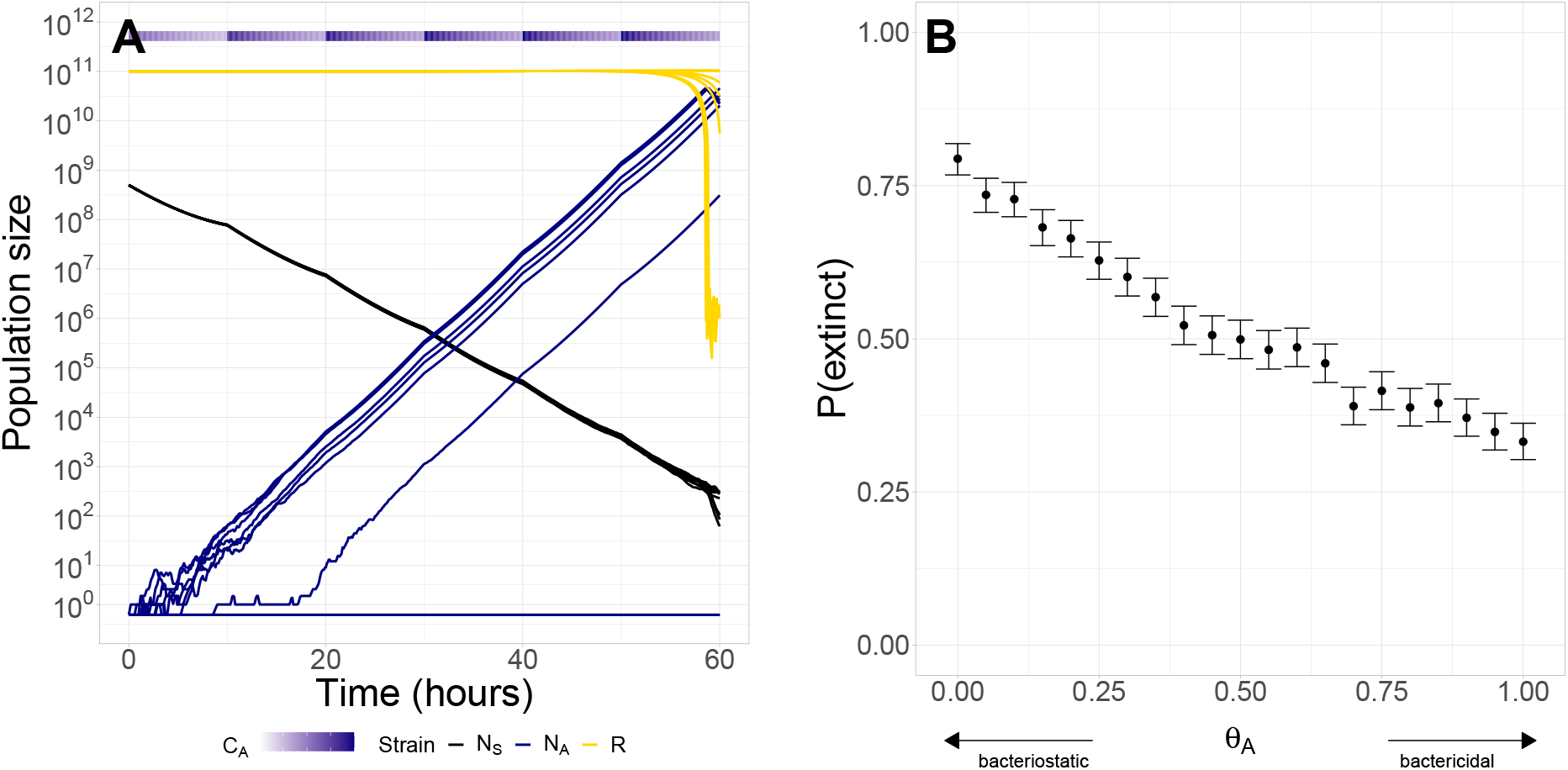
Effect of drug mode of action on resistance evolution, using monotherapy. A) shows example population dynamics in 10 simulation runs. B) shows the proportion of runs in which the population was not rescued by the resistant strain. 95% confidence intervals were generated using the standard error over 1000 simulation runs for each *θ*_*A*_ value. Parameter values were as in Table 1 except for *C*_*A*_(0) = 6, *C*_*B*_(0) = 0, *m*_*B*_ = 0.

Next, we extended the basic scenario to consider both cycling and combination therapy. Here, each drug varies independently on the bacteriostatic-bactericidal axis, so there are two degrees of freedom, with *θ*_*A*_ +*ϕ*_*A*_ = 1, *θ*_*B*_ +*ϕ*_*B*_ = 1. It is now not necessarily the case that the susceptible bacterial population will decline at the same rate across all combinations of modes of action, as bacteriostatic and bactericidal drugs act multiplicatively and additively, respectively, in our model. Again, we were interested in the proportion of simulation runs in which evolutionary rescue occurred.

For combination therapy, we found that one drug being fully bactericidal and the other being fully bacteriostatic maximized the probability of preventing AMR emerging (Figure 2A). This is because the same logic as in the monotherapy case applies as to why having at least one drug be bacteriostatic is beneficial. However, because there are diminishing returns to further bacteriostatic activity — the growth rate cannot be forced below zero — there is more benefit in the second drug being bactericidal, to rapidly reduce the susceptible population from *t* = 10 onwards.

**Figure 2:**
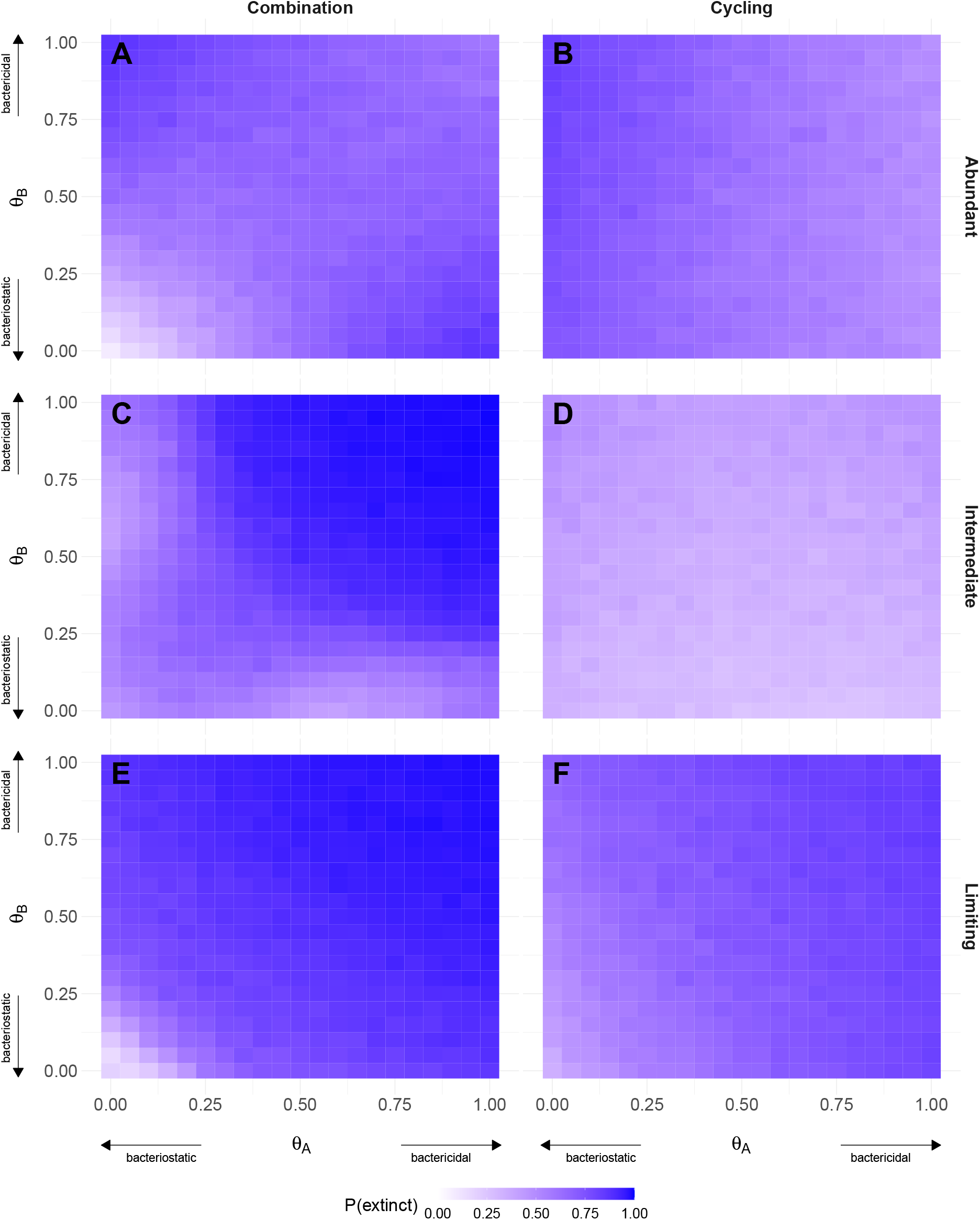
Effect of cycling and combination therapy, and varied resource constraints. Each panel shows how varying the mode of action of each drug changes the probability that the treatment successfully causes the bacterial population to go towards extinction, estimated over 1000 simulation runs for each grid square. The panels show each combination of resource availability (Abundant = A, B; Intermediate = C, D; Limiting = E, F) and therapy type (Combination = A, C, E; Cycling = B, D, F). Parameters used were the same as in Table 1 except for in combination treatments *S*(0) = 10^10^, *γ*_*j*_ = 0.35, *C*_*j*_(0) = 6, in resource-limiting treatments *R*(0) = 10^8^, *δ*_*i*_ = 0.25 and resource-intermediate treatments are halfway in between the two extremes with *R*(0) = 10^9.5^, *δ*_*i*_ = 0.325.

Similarly, for cycling therapy AMR is best averted by using a bacteriostatic drug *A* first and then a bactericidal drug *B* (Figure 2B). The diagonal symmetry of Figure 2A is broken here because the two drugs are applied at different times. It is now most important to first minimize replication, and hence chances for resistance mutations, by using a bacteriostatic drug *A*. The second drug *B* would then have less effect if it were also bacteriostatic, so it is preferable for it to be bactericidal.

There are no qualitative differences in these results when collateral sensitivity — increased susceptibility to one drug in cells resistant to the other — is included in the simulation (Figure S1). Increasing the elimination rate of the drugs from the patient does change the findings: because the concentration of the previous drug is then very low by the time the next drug dose is administered, the antagonism between two bacteriostatic drugs largely disappears, and now using two maximally bacteriostatic drugs is optimal (Figure S2).

All the results thus far have been using an initial resource concentration *R*(0) = 10^11^ that is in excess of what the population can consume during the time frame under consideration, making resource constraints negligible. If resources are less abundant, growth rates will be considerably constrained. As seen in Figure 2E,F binding resource constraints mean that the advantage of bacteriostatic drugs is reversed, and now AMR is best avoided by using two fully bactericidal drugs. This is because bacteriostatic drugs reduce growth rate proportionally, and so when resource constraints limit growth even in the absence of drugs, in absolute terms bacteriostatic drugs are less effective than their bactericidal counterparts. When resource constraints are noticeable but less severe, as seen in Figure 2C,D the effects are in between the more extreme scenarios, as expected.

Finally, if any resistant cells are already present when treatment is started, only the first drug that is used matters (in cycling therapy) and bactericidal drugs are more effective as they provide a greater chance of near-immediate death of the few resistant cells (Figure S3).

We also created an online interactive web-app version of the simulation: https://oscar-delaney. shinyapps.io/honours/. It has functionality significantly beyond that used in these analyses, which we hope will be useful for other researchers to quickly and visually explore AMR evolutionary dynamics.

## 4 Discussion

Our key finding — that bacteriostatic drugs are superior in monotherapy, and bacteriostatic and bactericidal drugs should be used together in combination or cycling therapy — is sensitive to the particular pharmacodynamic model used, and so our choice warrants further justification. There is no standard way to implement multi-drug bacteriostatic interactions, so we reasoned from first principles. To kill a cell, it is sufficient for any single essential cellular function to be disrupted, so it makes sense to model two bactericidal drugs targeting different cellular processes as acting additively: **D**_*i*_ = *δ*_*i*_ +*E*_*i*_(*C*_*A*_)*θ*_*A*_ +*E*_*i*_(*C*_*B*_)*θ*_*B*_ (known as Bliss independence in the literature [23]). However, the rate of cell division cannot be negative, so the total bacteriostatic activity of both drugs must be no more than unity, hence the Bliss independence method cannot simply be ported over to the bacteriostatic case. For a cell to divide, many separate physiological processes must function properly, and so the rate of division is proportional to the probability that each process is functional, which leads to our multiplicative model: **G**_*i*_ ∝ (1−*E*_*i*_(*C*_*A*_)*ϕ*_*A*_)(1−*E*_*i*_(*C*_*B*_)*ϕ*_*B*_). Thus, while there are many possible model configurations that could lead to different results, our implementation has theoretical support.

Like us, Nyhoegen and Uecker [16] hypothesise that “since replication and mutation are coupled, a drug-induced decrease in the replication rate will also reduce the per capita rate at which mutations occur” but they leave this speculation uninvestigated. Several theory papers have broached the relative merits of bactericidal and bacteriostatic antibiotics. Some make minimal distinctions between the two modes of action [24], while others define bacteriostatic drugs as less potent versions of bactericidal drugs, still affecting death rate rather than growth rate [25, 26]. Igler, Rolff, and Regoes [11] successfully implement bacteriostatic and bactericidal modes of action, and also speculate that bacteriostatic drugs may limit the evolution of AMR, however they do not investigate this, instead focusing on the effects of varying the pharmacokinetic model used. Newton, Ho, and Huang [27] also implement the difference between bactericidal and bacteriostatic drugs appropriately in their recent model, but they explore community dynamics among competing species, without considering the evolution of resistance.

Experimentalists have also investigated drug modes of action, finding that bacteriostatic and bactericidal drugs often interact antagonistically, putatively because bactericidal drugs work best when cells are dividing rapidly and uptaking many resources (and hence drug molecules) from the environment [28, 29]. Thus, if a bacteriostatic drug slows growth, the bactericidal drug may be less able to increase death rates. This differs from our result that one bactericidal and one bacteriostatic drug is optimal. The contrast arises from how bactericidal drugs are modelled: we assume that they act independently of growth rate, so relaxing this assumption to consider a death rate term proportional to the growth rate could improve the realism of our model. Another possible extension would be to consider pulsed as well as continuous resource supply, which has been shown in related contexts to alter population dynamics [30].

A meta-analysis with 9597 patients across 33 individual studies found no statistically significant difference between bacteriostatic and bactericidal drugs in terms of mortality or other individual patient outcomes [31]. However, due to the relative difficulty of collecting large-scale data on the evolution of resistance in clinical settings, we are aware of no studies comparing the proportion of patients that acquire resistance de novo when treated with bactericidal versus bacteriostatic antibiotics. While logistically challenging, this could be a valuable line of inquiry.

We hope that our theoretical analysis will motivate future experimental work to empirically test these hypotheses, to eventually inform clinical and public health decision-making for constraining AMR.

## Data availability

Additional figures are available in the supplementary file. All R code used for the simulation, generating the figures, and creating the web app, is available at https://github.com/Oscar-Delaney/ bacteriostatic.

