## Supplementary Figures for "Drug mode of action and resource constraints modulate antimicrobial resistance evolution"

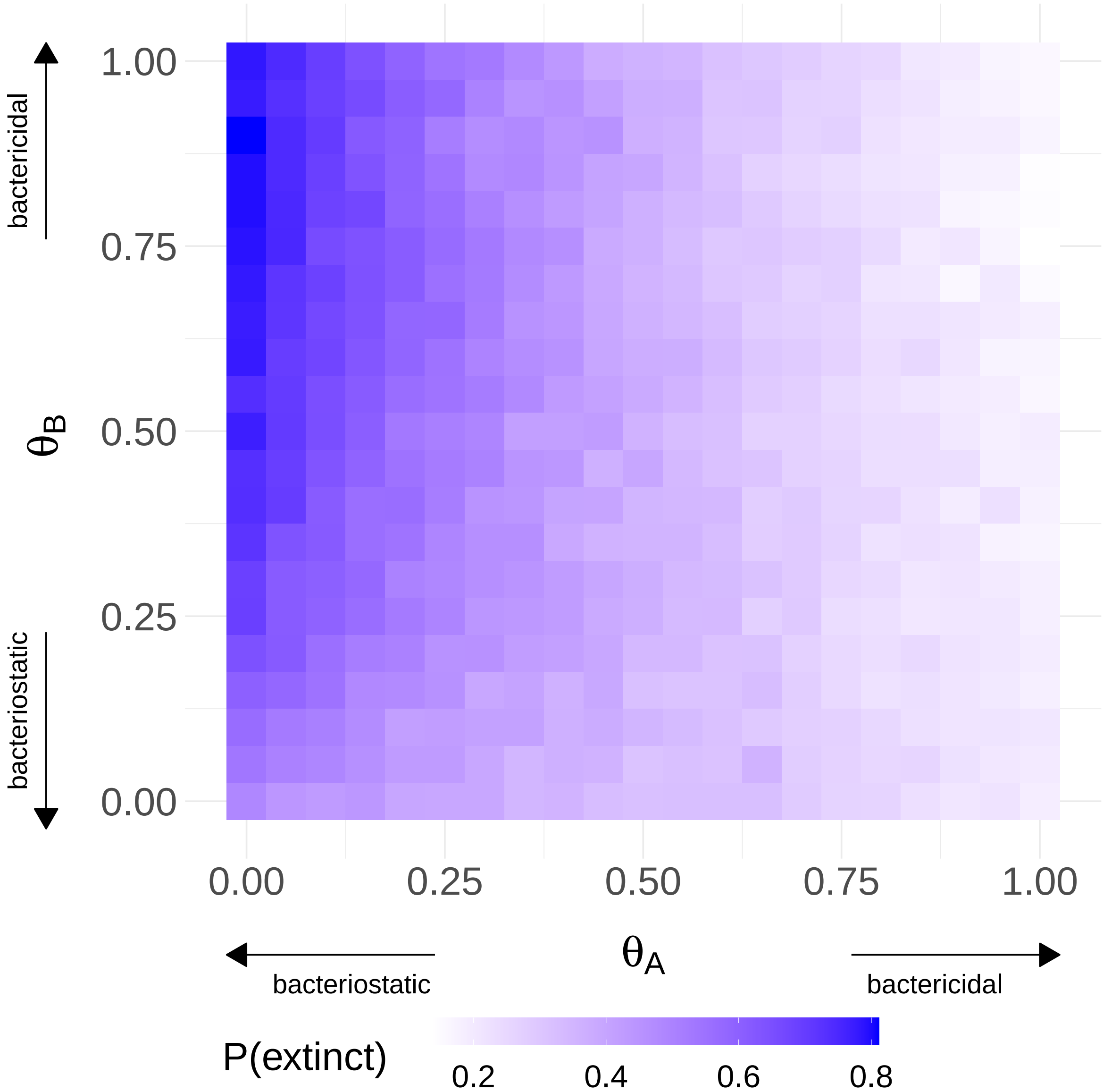


Figure S1: Under reciprocal collateral sensitivity resistance is best prevented by the first drug being bacteriostatic and the second drug bactericidal. Each grid square represents 1000 simulation runs. Cycling therapy with abundant resources was used, with parameter values the same as in Table 1 except $z_{B,A} = z_{A,B} = 0.5, \delta_i = 0.1, C_j(0) = 7$.

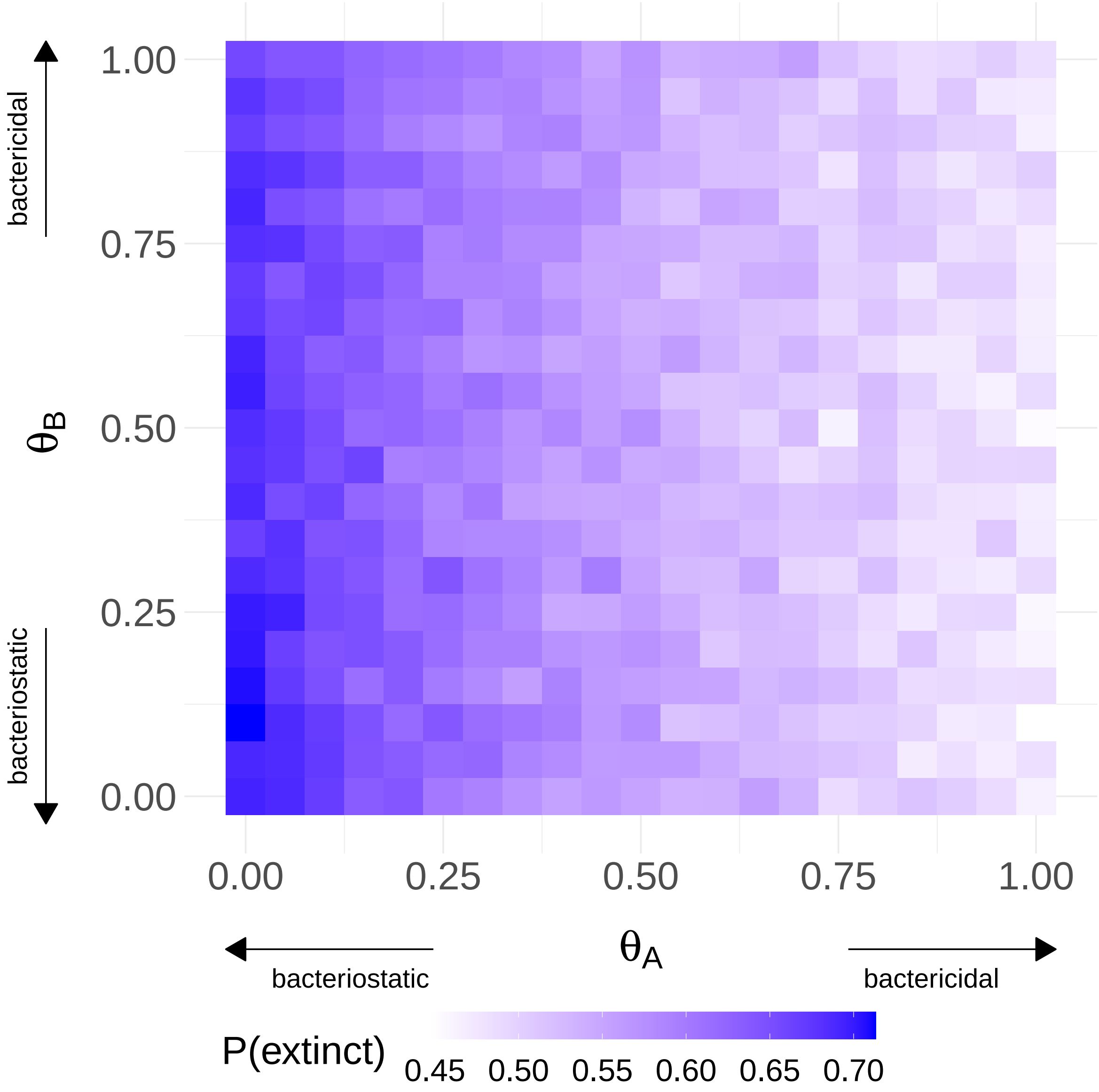


Figure S2: Faster drug elimination means both drugs can be bacteriostatic without encountering diminishing returns. Each grid square represents 1000 simulation runs. Cycling therapy with abundant resources was used, with parameter values the same as in Table 1 except $C_j(0) = 30, \delta_i = 0.45, \gamma_j = 0.5$.

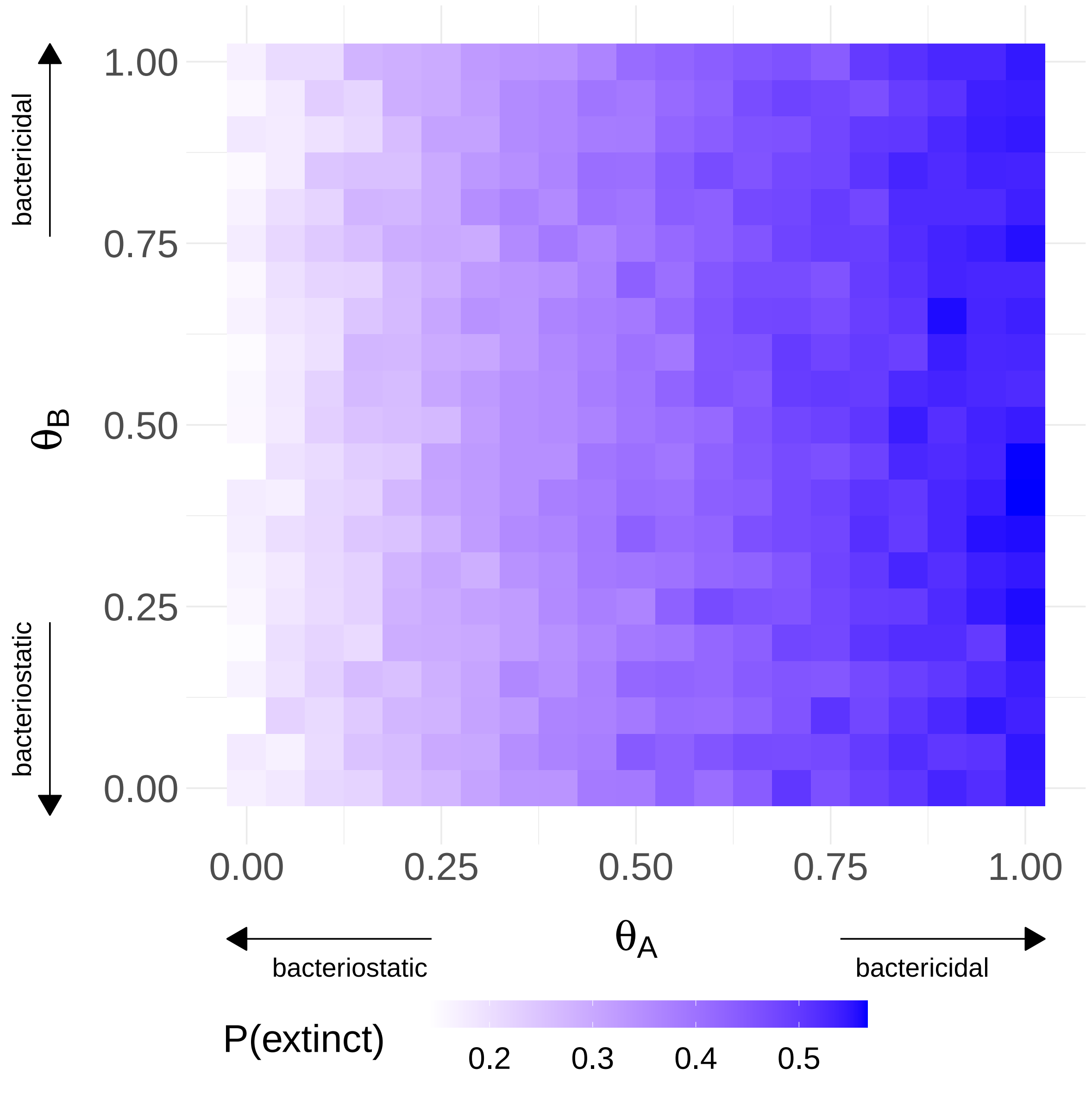


Figure S3: The presence of resistant cells prior to beginning treatment means that bactericidal drugs are more effective. Each grid square represents 1000 simulation runs. Cycling therapy with abundant resources was used, with parameter values the same as in Table 1 except $N_S(0) = 0, N_B(0) = 5, \delta_i = 0.25$.
